# Rich structure alphabets enable highest accuracy protein search

**DOI:** 10.64898/2026.07.24.740611

**Authors:** Robert C. Edgar

## Abstract

Protein structure databases have grown from thousands of experimentally determined structures to hundreds of millions of AI-predicted models, creating an urgent need for search methods that combine high accuracy with practical scalability. Here, I present the third generation of Reseek, a protein structure search algorithm achieving the highest overall accuracy (median rank 1) according to diverse metrics among tested methods including DALI, Foldseek and TM-align. Improved accuracy is obtained by parallel sequence alignment of many discrete alphabets capturing primary, secondary and tertiary features, giving a combined space of ***∼*10^22^** possible states. Separate statistical models are optimized for family, superfamily and fold discrimination, respectively, revealing distinct combinations of features that characterize each level. Hundreds of query structures can be searched a against a multimillion-structure database on a server computer in minutes, making large-scale structure search at state-of-the-art accuracy practical on commodity hardware.

## Introduction

Protein homology inference is a cornerstone of molecular biology, facilitating genome annotation, functional inference, phylogenetics and protein engineering. Structure is generally more conserved than sequence (Chothia and Lesk 1986), enabling identification of distant homologs by structure comparison. Recent advances in artificial intelligence (AI) have dramatically expanded the scale of structural biology. AlphaFold2 (Jumper et al. 2021) achieved accurate structure prediction from amino acid (aa) sequence and created the AlphaFold Protein Structure Database (Varadi et al. 2022) (AFDB, currently *∼*240M proteins). Subsequently, the ESMFold delivered faster structure prediction and enabled construction of the ESM Atlas database (Lin et al. 2023; Candido et al. 2026) which currently contains *∼* 1B structures, extending coverage deep into the metagenomic realm (Lin et al. 2023). This unprecedentedly rapid expansion from hundreds of thousands of solved structures to hundreds of millions of predicted structures has shifted the analysis bottleneck from structure inference to structure search, which canonically comprises construction and ranking of structure alignments. Previously popular structure alignment algorithms including DALI (Holm and Sander 1993) and TM-align (Zhang and Skolnick 2005) are computationally expensive and therefore do not scale to routine searches of AI-scale databases, prompting the development of Foldseek (Van Kempen et al. 2024) and Reseek (Edgar 2024) which can align a query structure to millions of proteins in seconds or minutes on a commodity computer. Foldseek and Reseek deploy broadly similar strategies based on representing structures as sequences of discrete states, thereby leveraging decades of research into computationally efficient algorithms for sequence alignment and search.

In this work, I present the third generation of Reseek with substantial improvements in search accuracy. Reseek (here, version 3 is understood unless otherwise stated) has highest rank by a diverse range of accuracy metrics while readily scaling to databases with millions or billions of structures. These improvements are achieved primarily through the development of a richer inventory of discrete structure alphabets and training of separate statistical models for inferring family, superfamily, and fold relationships. For each of these truth standards, two sets of default parameters are implemented: “fast” targeting higher speed at some cost in sensitivity and “sensitive” targeting higher sensitivity at some cost in speed.

Measuring search accuracy requires a trusted reference database. The SCOP40 subset of SCOP (Murzin et al. 1995) clustered at 40% aa identity is the *de facto* standard. SCOP classifies protein domains into families, folds and superfamilies. Superfamilies are intended to represent homologous groups. Folds are groups of superfamilies with similar conformation, which may be due to homology or convergence. Families are subgroups of superfamilies with unambiguous sequence homology. CATH (Orengo et al. 1997) provides a similar classification scheme with so-called topology playing a similar role to SCOP fold; CATH40 clustered at 40% aa identity provides an alternative reference together with SCOP40c (Edgar and Sahakyan 2025), a curated subset of SCOP40 which discards problematic short domains and several superfamilies assigned to different folds which are clearly homologous.

To compare search accuracy of different algorithms, I report several complementary types of plot: coverage vs. error (CVE) (Brenner et al. 1998), precision-recall (PR), and receiver-operator characteristic (ROC) (Davis and Goadrich 2006). Each of these plots is constructed twice, using distinct strategies designed to model distinct scenarios for search: “collection”, finding many high-confidence homologs, and “categorization”, classifying the query into a category annotated by the search database (e.g., fold). For categorization, only the top hit is considered and challenging searches are modeled using a leave-category-out strategy. For example, to model classification to superfamily other members of the same family are left out, and for fold classification other members of the same superfamily are left out. Algorithm rankings are calculated using summary metrics derived from these plots.

## Results

### Reseek algorithm

The algorithm workflow has three stages (Fig. 1): a *k*-mer search for 2-hit diagonals (*κ* filter), an SIMD-accelerated Smith-Waterman (Smith et al. 1981) step (*ν* filter) which discards low-scoring alignments, and a final Smith-Waterman alignment using a compound alphabet called Mega*^′^* (*M ^′^*). Kappa (*κ*) is a 32-state alphabet designed to facilitate identification of similar *k*-mers. Nu (*ν*) is a 256-state alphabet designed to capture rich information while facilitating SIMD optimizations and compact database storage, both of which benefit from representing a residue state in a single byte. *M ^′^* is a weighted product over many smaller alphabets giving *∼* 5 *×* 10^22^ possible states which integrate sequence and structure information captured by all implemented alphabets. *M ^′^* profiles and *M ^′^* alignments are constructed on-the-fly and are relatively time-consuming to compute, but this time is typically less than half of the total because a large majority of query-database pairs are eliminated by the filtering steps. If the *M ^′^* alignment has high score, the reversed *M ^′^* profile is also aligned to the database profile. A test statistic (TS) is calculated by a weighted combination of the forward and reverse *M ^′^* scores plus LDDT (Mariani et al. 2013) and DALI scores implied by the forward alignment. A *P*-value is calculated from TS using log-linear fitting according to the desired truth standard (family, superfamily or fold) (Edgar and Sahakyan 2025).

**Fig. 1.**
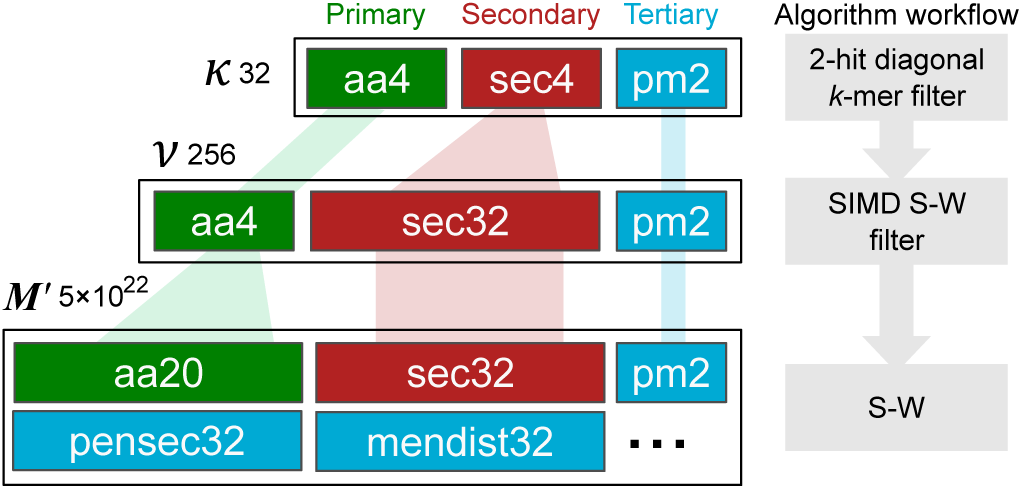
Workflow of the Reseek algorithm. First is a 2-hit diagonal filter using Kappa alphabet sequences. Second is SIMD-accelerated Smith-Waterman of Nu sequences of the query and reversed query (if the forward *ν* score is high enough). Third is Smith-Waterman of *M ^′^* profiles, including the reversed query if the forward *M ^′^* score is high enough. Finally LDDT and DALI scores implied by the forward *M ^′^* algorithm are used together with alignment scores to calculate the test statistic according to the chosen model.

### Search accuracy

Fig. 2 reports CVE, PR and ROC plots using SCOP40c as reference and superfamily as truth standard, showing that Reseek has substantially higher accuracy than the other tested methods in the high-scoring regime. In this plot, high-scoring is operationally defined by the axis ranges as up to 10 errors per query (corresponding to *E*-value *≤* 10), precision *≥* 70%, and FPR *≤* 10*^−^*^4^, respectively. Corresponding plots for other references and truth standards are provided in supplementary material.

**Fig. 2.**
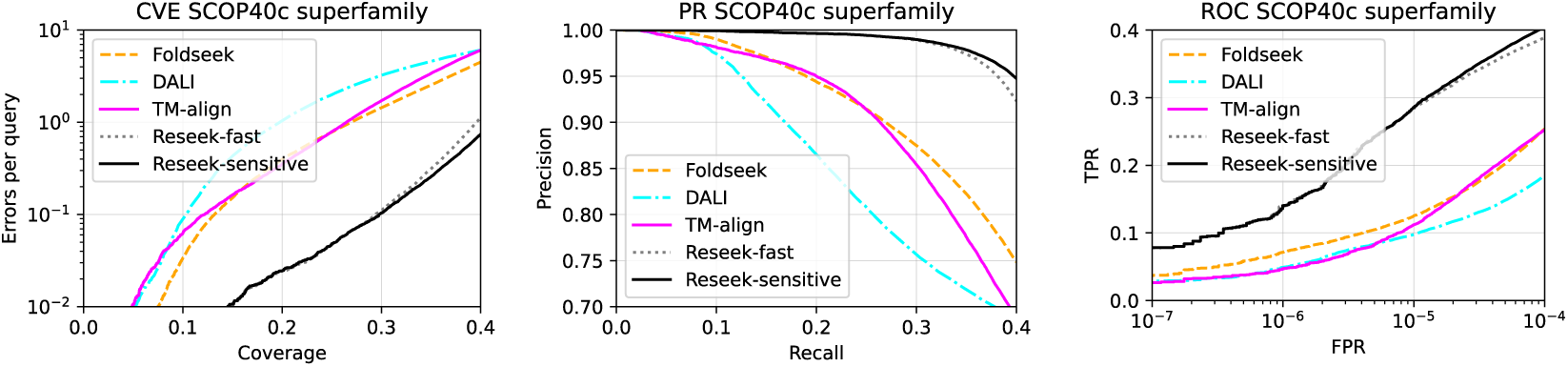
Accuracy plots for SCOP40c superfamily. The figure gives CVE, PR and ROC plots for the tested algorithm using SCOP40c as reference and superfamily as truth standard. For a CVE plot, a lower curve is better while for PR and ROC a higher curve is better. These curves show that Reseek-sensitive has higher accuracy than other tested methods in the high-scoring regime (up to 10 errors per query for CVE, *≥* 70% precision for PR and *<* 10*^−^*^4^ FPR for ROC).

### Method rankings

Table 3 reports method ranks by each of the considered reference, truth standards, and metrics. Most accurate is Reseek-sensitive, with median rank 1, followed by Reseek-fast with median rank 2. Table 4 reports pair-wise *P*-values according to the Wilcoxon signed-rank test, showing that the higher rank of Reseek-sensitive has significance ranging from *P* = 5.4 *×* 10*^−^*^4^ against TM-align to 1.1 *×* 10*^−^*^5^ against Foldseek.

### Model weights

Figs. 3 and 4 show alphabet weights in *M ^′^* and term weights in TS, respectively, which were tuned by Hooke-Jeeves optimization (Hooke and Jeeves 1961). The optimized weights exhibit an unexpectedly sharp partitioning of feature importance which can be interpreted as reflecting essential differences between family, superfamily and fold categories. The contribution of primary sequence (aa20 and aa4) drops from 61% aa20 for family to 18% aa4 for fold, while the weight of nensec32 (nearest-neighbor secondary structure) increases from 3.6% for family and 1.6% for superfamily to 27% for fold, where it is the highest-weighted alphabet. (Alphabets with zero weight, including aa4 for family and superfamily, and aa20 for fold, are not shown). The weight of sec32 is 4.5% family, 5.9% superfamily and 10% fold. Overall, the progression is decreasing weight for primary sequence and increasing weight for secondary and tertiary structure features. In particular, fold strongly emphasizes secondary structure of a residue and its nearest neighbor. Remarkably, automated optimization almost completely separated the coordinate-based terms by SCOP level: LDDT contributed almost exclusively to family discrimination, whereas DALI contributed almost exclusively to fold discrimination (Fig. 4). LDDT was designed to compare predicted and experimentally determined structures, considering only C*α* pairs within a local spatial neighborhood (*≤* 15 Å), whereas DALI was designed to recognize topological similarity by employing a smooth distance-dependent weighting with a characteristic length scale of 20 Å (*α* parameter in (Holm and Sander 1993)), allowing longer-range distances to contribute with decreasing weight (Fig. 9). Taken together, these results suggest that families are largely discriminated by amino acid sequence similarity and spatially adjacent structure reflected by LDDT; superfamilies are discriminated by log-odds alignment scoring while C*α* coordinates contribute little; and folds are discriminated by spatially adjacent secondary structures and long-distance coordinates captured by the DALI score. The remaining contribution of primary sequence to fold discrimination may be explained by composition bias in secondary structure (Chou and Fasman 1974), and therefore does not necessarily imply that sequence homology is playing a role. Noting that log-odds scoring implicitly models relatedness via substitutions, these results are consistent with a hypothesis that superfamilies correspond well to homologous groups while different superfamilies in the same fold tend to be convergent.

**Fig. 3.**
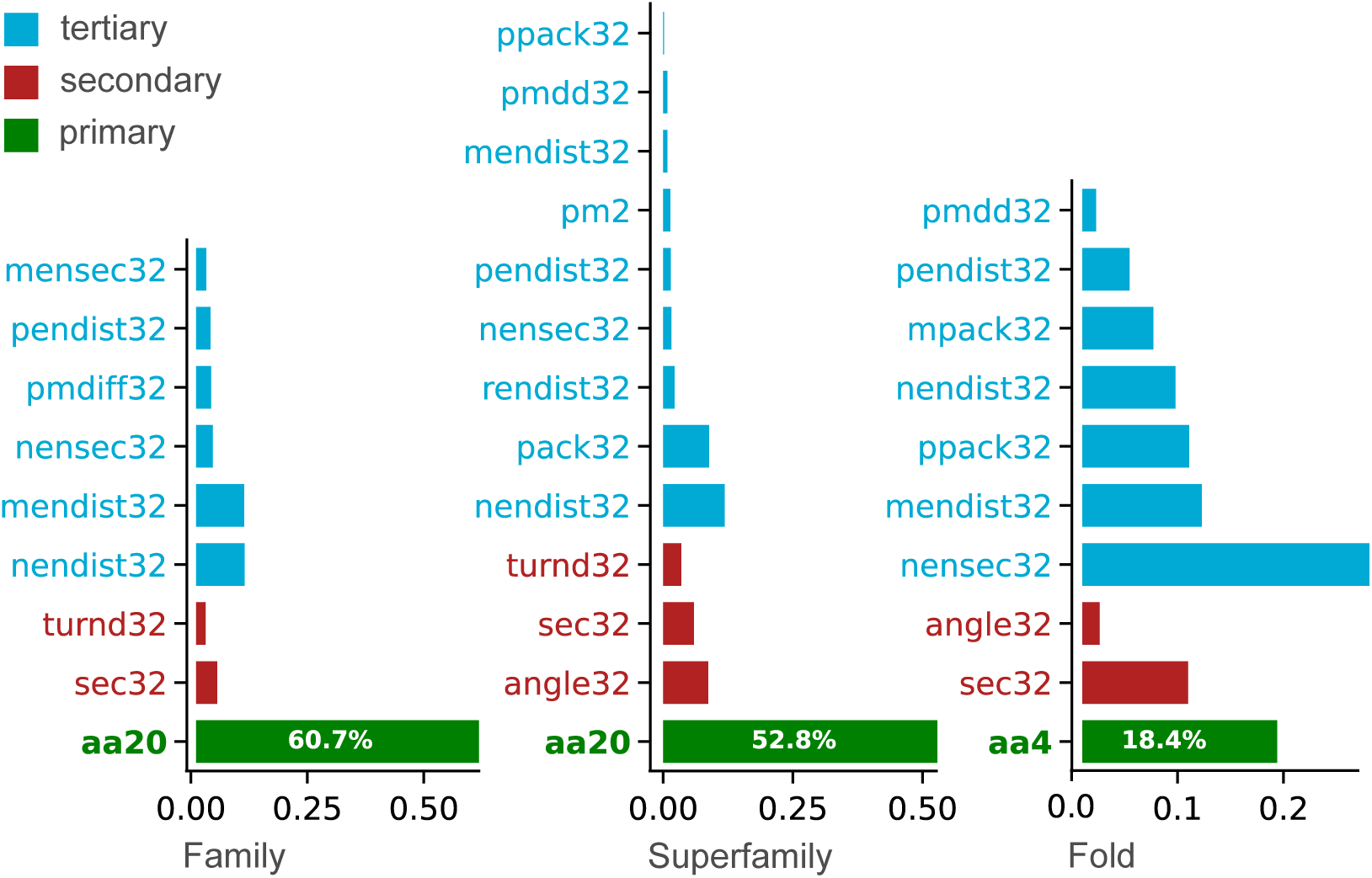
Alphabet weights in default statistical models. Weights are trained to discriminate family, superfamily and fold, respectively, and are normalized to sum to 1. Note that the contribution of amino acid similarity decreases while secondary and tertiary structure weights increase from family to fold.

**Fig. 4.**
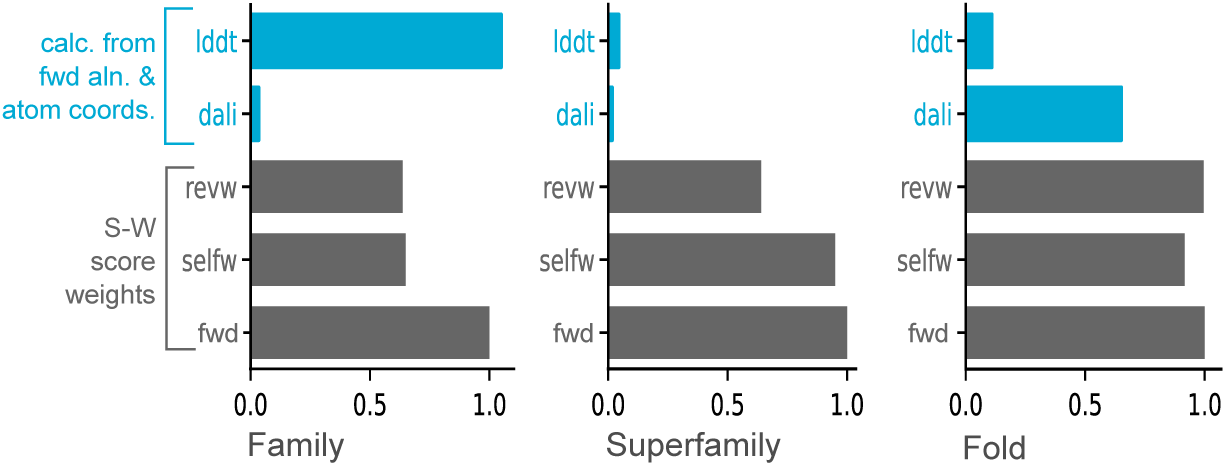
Test statistic component weights. The test statistic (TS) is a weighted sum of LDDT, DALI, *revw* (*M ^′^* score of the reversed query against the target), *revw* (sum of the *M ^′^* score of the query against its reversed profile and the target against its reversed profile), and *fwd* (the *M ^′^* score of the query and target). By convention, *fwd*=1. Remarkably, this shows that automated optimization almost completely separated the coordinate-based terms by level. For superfamily, the coordinate-based weights LDDT and DALI are low, leaving log-odds terms which model homology via substitutions. LDDT, which focuses on shorter distances, contributes almost exclusively to family discrimination, whereas DALI, which considers longer distances, contributes almost exclusively to fold discrimination (see also Fig. 9).

### Scaling to large databases

Reseek stores a so-called inverted *k*-mer index on the query in RAM, i.e. an index mapping *k*-mers to query structures. By contrast, Foldseek stores an inverted index of database *k*-mers in RAM. These designs give quite different scaling behavior, with time and memory trade-offs varying strongly with query size, database size, available RAM and hit rate (i.e., the fraction of the database which has detectable similarity to the query set). Most obviously, Foldseek requires RAM which scales roughly linearly in database size, while Reseek requires RAM which scales roughly linearly in query size but does not increase with database size. Each method may achieve sub-linear time scaling in query and/or database size, depending on the data, and each method may benefit (or not) from splitting the query into chunks which are processed separately, or from splitting the database into chunks. Given these widely different characteristics, comparative validation of scaling using commensurate metrics is ruled out. Here, to demonstrate that both methods are practical at large scale, I report two examples of searching a database with tens of millions of structures, noting that the results should be considered anecdotal rather than a rigorous comparison. Foldseek and Reseek were run against AFDB50 (*∼* 54M structures) using 100 and 1,000 randomly-chosen structures from the Protein Data Bank (Berman et al. 2000) as queries. CPU was Intel Xeon 6978P processor (240 cores) with 768GB RAM, specifying 64 threads on the command lines. As seen in Table 1, elapsed times were on the order of a few minutes to a few hours, while memory requirements ranged from 4.4Gb (Reseek-fast with query size 100) to 284Gb (Foldseek with query size 1,000).

**Table 1.** Computational cost of large database search. Foldseek and Reseek were run against AFDB50 (*∼* 54M structures) using 100 and 1,000 randomly-chosen solved structures as queries. CPU was Intel Xeon 6978P processor (240 cores) with 768GB RAM, specifying 64 threads on the command lines.

| Query<br>structs | Elapsed time |  |  |
| --- | --- | --- | --- |
|  | Foldseek | Reseek-fast | Reseek-sensitive |
| 100 | 9m | 6m | 23m |
| 1000 | 19m | 61m | 4h |

Table 1. Computational cost of large database search. Foldseek and Reseek were run against AFDB50 ( $\sim 54$ M structures) using 100 and 1,000 randomly-chosen solved structures as queries. CPU was Intel Xeon 6978P processor (240 cores) with 768GB RAM, specifying 64 threads on the command lines.
| Query<br>structs | Max memory |  |  |
| --- | --- | --- | --- |
|  | Foldseek | Reseek-fast | Reseek-sensitive |
| 100 | 227Gb | 4.4Gb | 4.5Gb |
| 1000 | 284Gb | 6.6Gb | 6.7Gb |

### Hit sorting by statistical models

Reseek currently supports three statistical models by default, which were trained to discriminate family, superfamily and fold categories, respectively. This novel design raises the question of how well a model sorts hits when evaluated by the “wrong” truth standard. This is addressed in Fig. 5, which shows CVE plots using SCOP40c as reference for all nine combinations of (training standard, evaluation standard), which shows that there are substantial differences in sort order when different models are used. Presumably, this reflects emphasis on different features such as local conformation for family vs. large-scale secondary structure organization for fold. Therefore, it is important for a user to choose the model which corresponds best to the objectives of their search.

**Fig. 5.**
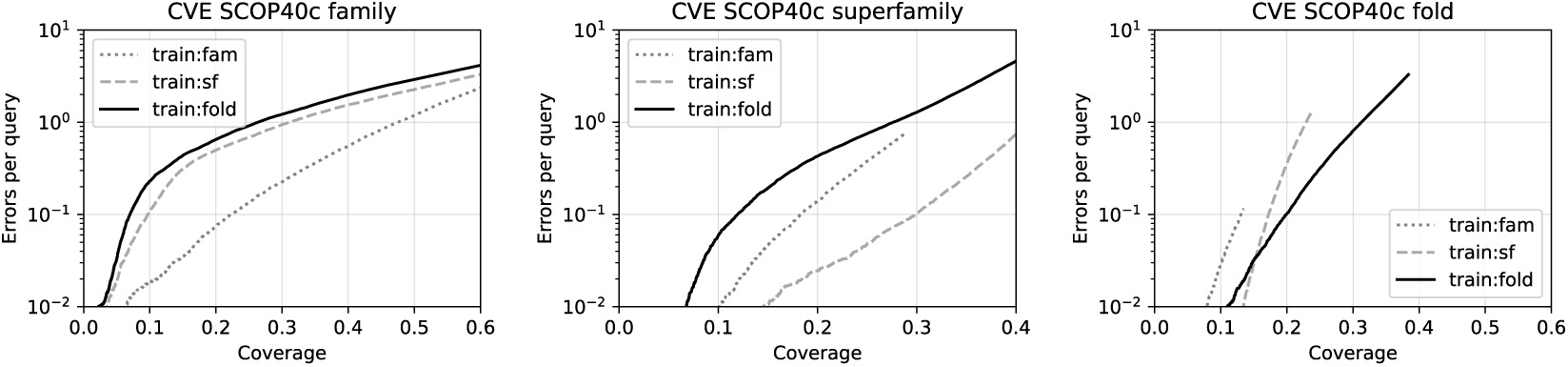
Divergent hit sorting by statistical models. The figure shows CVE plots using SCOP40c as reference for all nine combinations of (training standard, evaluation standard) for truth standards (family, superfamily, fold), showing that there are substantial differences in sort order when different models are used. Some curves are truncated at the higher error / higher coverage end due to the default *P*-value threshold (0.001).

## Methods

### Alphabet construction

Table 2 summarizes alphabets implemented in *M ^′^*. In common with most other structure alignment and search algorithms, Reseek considers C*α* atom coordinates only, which are conventionally specified as (*x, y, z*) coordinates. Axes are chosen arbitrarily, in the sense that a different choice of axes corresponds to a rigid translation and/or rotation, giving an equivalent set of coordinates. Distances between C*α* atoms can be viewed as inherent to a structure, while their (*x, y, z*) coordinates require adding a frame of reference, and it is therefore is natural and convenient to convert coordinates into a so-called contact map (CM), i.e. an all-vs-all distance matrix (Fig. 6). A CM is unchanged under translation and rotation, and a CM suffices to construct (*x, y, z*) coordinates if needed (Torgerson 1952). Consider a given residue *X* (the *focus* residue). An alphabet *A* is specified by a function *R_A_*(*X,* CM) which yields an integer in the range 1*…|A|*, where *|A|* is the alphabet size. *R_A_* must be independent of the axis choice, and without loss of generality the input to *R* can therefore be defined as the contact map and position of the focus residue. Conceptually, it can be helpful to divide the contact map into regimes corresponding to secondary structure, intra-domain tertiary structure, and inter-domain tertiary structure (Fig. 6). Secondary structure corresponds to local conformation such as alpha helix or beta sheet, which can be recognized from all-vs-all distances in a window of *± ∼* 3 positions around *X* (Levitt and Greer 1977), i.e. the CM of a *∼* 7-mer centered on *X*. Tertiary structure within a domain can be loosely described by interactions between residues which are not adjacent in the primary sequence. Domains are typically *∼* 100 .. 200*aa*, with few domains longer than *∼* 500*aa*; interactions between residues at longer sequence offsets therefore tend to be between different domains.

**Fig. 6.**
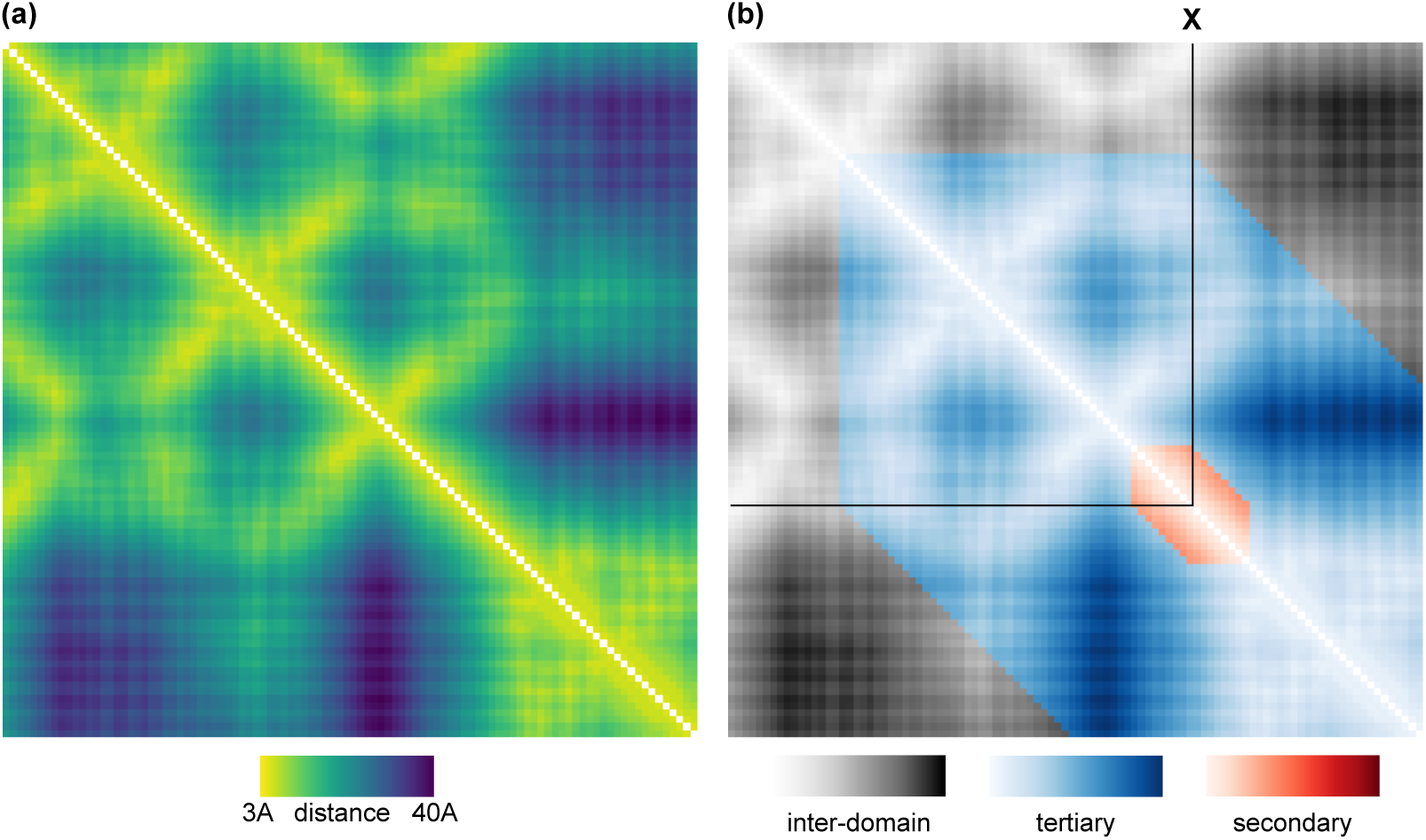
Contact map. Panel (a) shows a representative contact map, i.e. all-vs-all distance matrix between C-alpha atoms, constructed from the first 200 residues in 1hhs chain A. Distances are represented on a color scale ranging from 3A to 40A (clipped). Panel (b) divides the map for focus residue *X* into three conceptual regimes: secondary structure (all-vs-all distances for a 7-mer centered on *X*), intra-domain tertiary structure (distances up to the end of the domain at offsets of around 100 to 200 residues from *X*), and inter-domain tertiary structure (at longer offsets from *X*).

**Table 2.** Component alphabets implemented in Mega*^′^*. *Name* is alphabet name including size, *D/Q* is *D* if the alphabet is inherently discrete or *Q* if the alphabet is “quantized”, i.e. derived from a real-valued feature by binning.

| Category | Name | D/Q | Description |
| --- | --- | --- | --- |
| Primary sequence | aa20 | D | Standard 20-letter aa alphabet |
|  | aa4 | D | Reduced aa {ACHPST}, {DEKNQR}, {FILMVWY}, {G} |
| Neighbor distance | nendist32 | Q | Nearest neighbor distance $d(\mathbf{X}, \text{NEN})$ |
| | rendist32 | Q | Reverse neighbor distance $d(\mathbf{X}, \text{REN})$ |
| | pendist32 | Q | Plus neighbor distance $d(\mathbf{X}, \text{PEN})$ |
| | mendist32 | Q | Minus neighbor distance, $d(\mathbf{X}, \text{MEN})$ |
| | pm2 | D | 1 if $d(\mathbf{X}, \text{PEN}) \geq d(\mathbf{X}, \text{NEN})$, otherwise 0 |
| | pmdd32 | Q | $d(\mathbf{X}, \text{PEN}) - d(\mathbf{X}, \text{NEN})$, clipped at 20Å |
| | pmdiff32 | Q | $d(\mathbf{X}, \text{PEN}) - d(\mathbf{X}, \text{NEN})$, clipped at 40Å |
| Secondary structure | sec32 | D | Secondary structure of $\mathbf{X}$, see Supp. Fig. TODO |
|  | sec4 | D | Reduced sec32 {AB}, {CDGINZ}, {EFJLKMORSTacdef}, {HPQUVWXYb} |
|  | nensec32 | D | Nearest neighbor secondary structure, i.e. sec32 of NEN |
|  | rensec32 | D | Reverse neighbor secondary structure, i.e. sec32 of REN |
|  | pensec32 | D | Plus neighbor secondary structure, i.e. sec32 of PEN |
|  | mensec32 | D | Minus neighbor secondary structure, i.e. sec32 of MEN |
| | angle32 | Q | Angle subtended by Cα atom triangle ($\mathbf{X}-4, \mathbf{X}, \mathbf{X}+4$) |
| | turnd32 | Q | Distance $d(\mathbf{X}-5, \mathbf{X}+5)$, small if $\mathbf{X}$ is in a turn |
| Packing | pack32 | Q | Number of Cα atoms at distance $\leq 15$ Å from $\mathbf{X}$ within window $+/-12...256$ residues in primary sequence |
| | ppack32 | Q | Ås pack except window is plus only i.e. $+12...256$ |
| | mpack32 | Q | Ås pack except window is minus only i.e. $-12...256$ |
| Compound | Kappa32 ($\kappa$) | $n/a$ | $\text{aa4} \times \text{sec4} \times \text{pm2}$ |
| | Nu256 ($\nu$) | $n/a$ | $\text{aa4} \times \text{sec32} \times \text{pm2}$ |

**Table 3.** Algorithm ranks. For each summary metric, the table lists ranks for each of the tested algorithms. A metric is specified as Type*>*Truth*>*Reference. Types Sum3 and PR90 summarize “collection” accuracy from CVE and PR, respectively. Top3 summarizes “categorization” accuracy where the top hit above the cutoff is used to annotate the query. The median ranks for Reseek-sensitive and Reseek-fast are 1 and 2, respectively. TM-align and DALI were not tested on CATH40 due to high computational cost.

| Metric | DALI | Foldseek | TM-align | Reseek-fast | Reseek-sens |
| --- | --- | --- | --- | --- | --- |
| Sum3>family>scop40 | 5 | 4 | 2 | 3 | 1 |
| Sum3>family>scop40x | 5 | 4 | 3 | 2 | 1 |
| Sum3>fold>scop40 | 4 | 5 | 2 | 3 | 1 |
| Sum3>fold>scop40x | 1 | 5 | 2 | 4 | 3 |
| Sum3>superfamily>cath40 |  | 3 |  | 2 | 1 |
| Sum3>superfamily>scop40 | 5 | 4 | 3 | 2 | 1 |
| Sum3>superfamily>scop40x | 5 | 3 | 4 | 2 | 1 |
| Sum3>superfamilyx>cath40 |  | 3 |  | 2 | 1 |
| Sum3>superfamilyx>scop40 | 4 | 5 | 3 | 2 | 1 |
| Sum3>superfamilyx>scop40x | 1 | 5 | 4 | 3 | 2 |
| PR90>family>cath40 |  | 1 |  | 3 | 2 |
| PR90>family>scop40 | 1 | 2 | 2 | 2 | 2 |
| PR90>family>scop40x | 5 | 4 | 3 | 2 | 1 |
| PR90>fold>cath40 |  | 1 |  | 3 | 2 |
| PR90>fold>scop40 | 4 | 5 | 2 | 3 | 1 |
| PR90>fold>scop40x | 3 | 5 | 1 | 4 | 2 |
| PR90>superfamily>cath40 |  | 3 |  | 2 | 1 |
| PR90>superfamily>scop40 | 5 | 4 | 3 | 2 | 1 |
| PR90>superfamily>scop40x | 5 | 3 | 4 | 2 | 1 |
| PR90>superfamilyx>cath40 |  | 1 |  | 3 | 2 |
| PR90>superfamilyx>scop40 | 3 | 5 | 4 | 2 | 1 |
| PR90>superfamilyx>scop40x | 1 | 5 | 2 | 4 | 3 |
| Top3>topfold>cath40 |  | 3 |  | 2 | 1 |
| Top3>topfold>scop40 | 4 | 5 | 3 | 2 | 1 |
| Top3>topfold>scop40x | 5 | 4 | 3 | 2 | 1 |
| Top3>topsf>scop40 | 5 | 4 | 3 | 2 | 1 |
| Top3>topsf>scop40x | 4 | 3 | 5 | 1 | 2 |
| Sum3>family>cath40 |  | 1 |  | 3 | 2 |
| Sum3>fold>cath40 |  | 3 |  | 2 | 1 |
| Median | 4 | 4 | 3 | 2 | 1 |
|  | DALI | Foldseek | TM-align | Reseek-fast | Reseek-sens |

**Table 4.** Pair-wise algorithm comparisons. This table gives *P*-values for Wilcoxon test on the ranks in Table 3, ‘*>*’ indicates that the algorithm in the row is more accurate than the algorithm in the column with *P <* 0.05, ‘*∼*’ for (Foldseek,DALI) and (Reseek-fast,TM-align) indicates that the difference is not significant. These results show that Reseek-sensitive is more accurate than each of the other tested algorithms with *P*-value ranging from 0.0007 (DALI) to 2.6 *×* 10*^−^*^6^ (Foldseek).

|  |  |  |  |  |
| --- | --- | --- | --- | --- |
| **Foldseek** | **~0.4991** |  |  |  |
| **TM-align** | **>0.02211** | **>0.003588** |  |  |
| **Reseek-fast** | **>0.01988** | **>0.002765** | **~0.2619** |  |
| **Reseek-sensitive** | **>0.0007144** | **>1.198e-05** | **>0.0005855** | **>2.705e-06** |
|  | **DALI** | **Foldseek** | **TM-align** | **Reseek-fast** |

Reseek characterizes secondary structure by considering a sparse subset, CM_7_(*X*), of the CM for the 7-mer centered on *X* (Fig. 7). CM_7_(*X*) is assigned to one of 32 representatives identified by *K*-means clustering of all 7-mers in SCOP40, setting *K* = 32. This alphabet is called sec32.

**Fig. 7.**
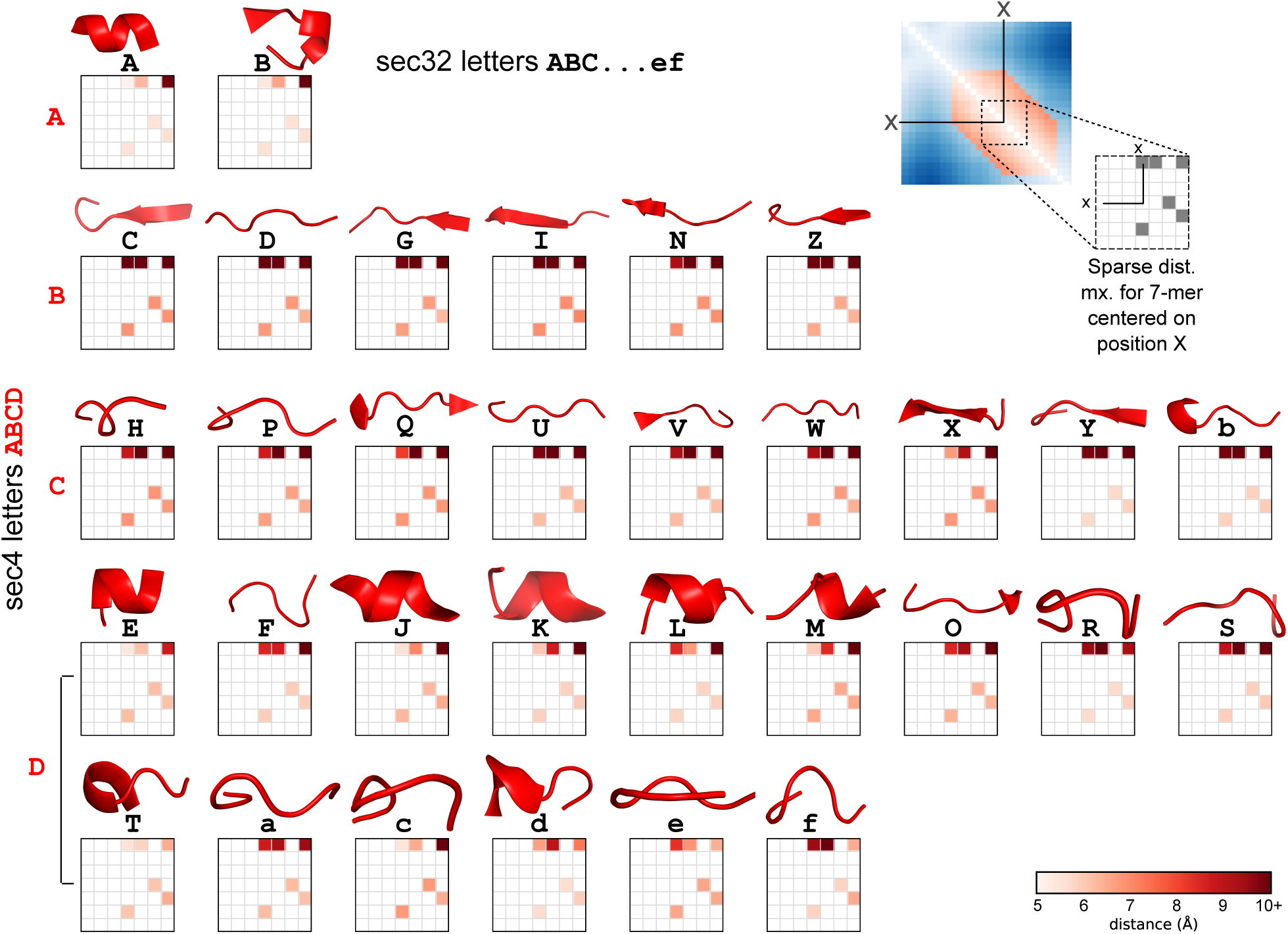
Secondary structure alphabets. The sec32 alphabet was constructed by *K*-means clustering (*K*=32) of sparse all-vs-all 7-mers in SCOP40, giving 32 representative 7*×*7 matrices with 6 cells filled. The sec32 letter for residue *X* is assigned by finding the best-matching representative. A letter in the reduced alphabet sec4 has 1:*n* correspondence with sec32 letters, enabling near-instantaneous conversion of a letter in sec32 or *ν* to a letter in sec4 (note that sec32 is a component of *ν*).

Interactions can be characterized by identifying “Euclidean neighbors”, i.e. residues which are closest in 3D space but not adjacent in the sequence. In Reseek, Euclidean neighbors must be separated by a minimum of 12 and maximum of 256 positions in the primary sequence. Four neighbors are defined: Nearest, Reverse, Plus and Minus (NEN, REN, PEN and MEN, see Fig. 8). Two pairs of these are identical: either NEN=PEN and REN=MEM, or NEN=MEN and REN=PEN, depending on whether the nearest Euclidean neighbor is on the C- or N-terminal side of *X*. A given interaction is characterized by the secondary structure of the neighbor, for example ne`nsec32` is the `sec32` letter for NEN(*X*), and the distance to the neighbor, for example `nendist32` is the distance to NEN(*X*), divided into 32 bins by specifying 31 thresholds that partition the distribution into roughly equal-sized ranges (see (Edgar 2024) for discussion of this approach to discretizing a real-valued feature).

**Fig. 8.**
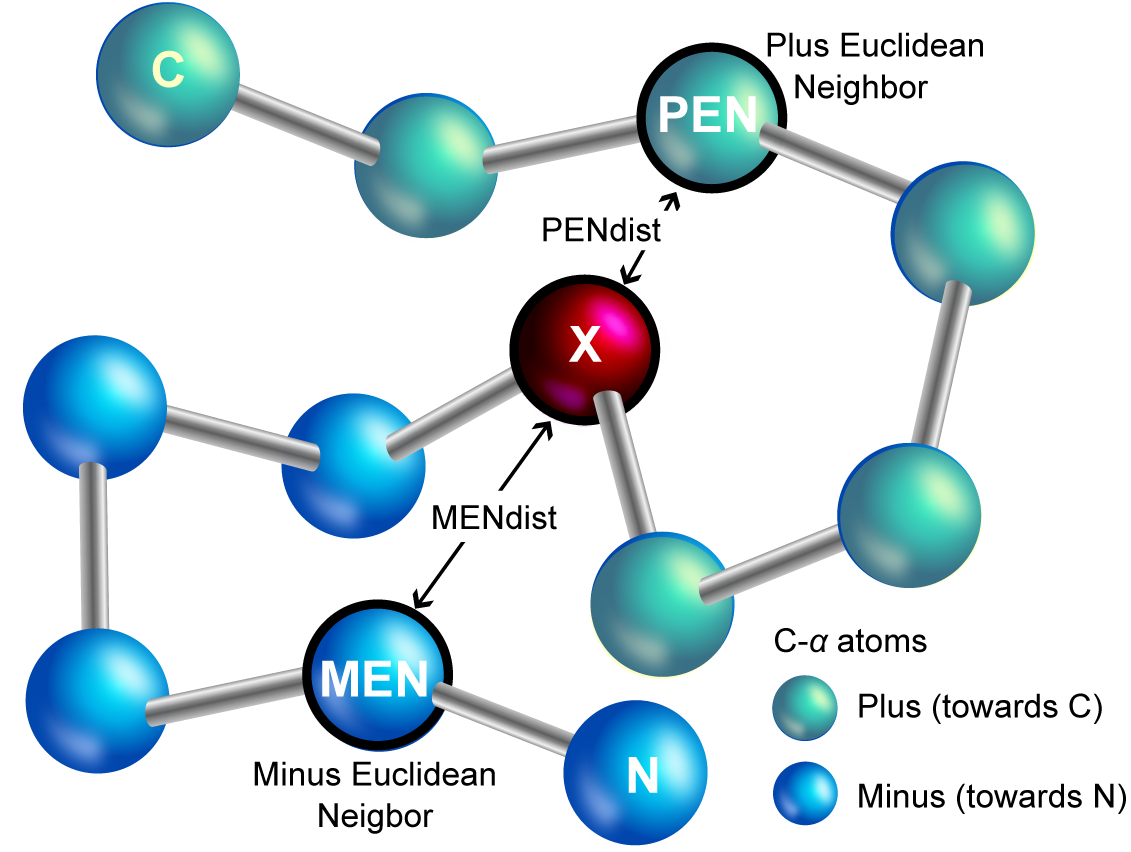
The focus residue and its Euclidean neighbors. The structure state of a given residue (the focus residue, *X*) is partly characterized by identifying its nearest Euclidean neighbors, i.e. the closest residues in 3D space (excluding residues immediately adjacent in the chain). The Plus Euclidean neighbor (PEN) is the closest residue in the C-terminal direction, while the Minus Euclidean neighbor (NEN) is closest in the N-terminal direction. The Nearest Euclidean Neighbor (NEN) is the closer of (PEN, NEN) and the Reverse Euclidean Neighbor is the further of (PEN, NEN). Distance to a neighbor is characterized by binning, e.g. mendist32 is the distance to MEN divided into 32 bins by setting 31 thresholds. The secondary structure of a neighbor is its sec32 letter (see Fig. 7).

**Fig. 9.**
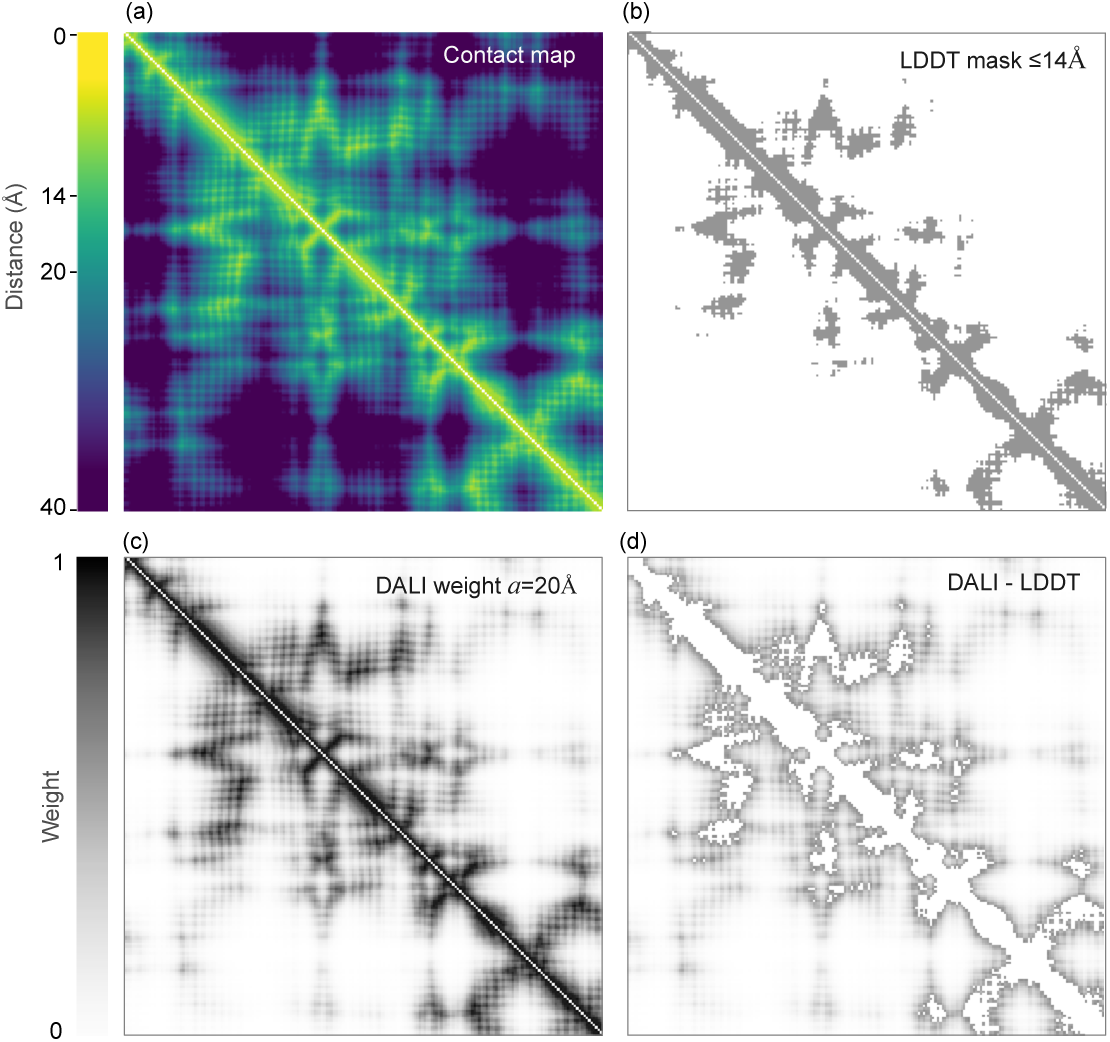
LDDT and DALI emphasize distinct distance scales. (a) Contact map for the first 400 residues in 5awa. (b) Distances *≤*14A considered by LDDT are shown as gray, longer distances are ignored by LDDT and are colored white. (c) Distances colored according to their DALI weight factor, which has a characteristic length scale *α*=20. Distances *>α* are included, but with lower weight. (d) As (c), but with distances considered by LDDT colored white. Thus, gray colors in (d) show distances which are considered by DALI but not by LDDT. This illustrates that LDDT focuses on shorter-scale conformation while DALI considers longer-range relationships. See also Fig. 4.

Kappa32 (*κ*) is the compound alphabet `aa4 × sec4 × pm2`, and Nu256 (*ν*) is `aa4 × sec32 × pm2`. This gives a 1 : *n* relationship between a *κ* letter and *ν* letters, enabling near-instantaneous calculation of a *κ* sequence from the *ν* sequence which is stored in database files.

### Algorithm workflow

The Reseek algorithm workflow is shown in Fig. 1. First stage is a 2-hit diagonal filter (Altschul et al. 1997) using *k*-mers in the *κ* alphabet where *k* = 7 and 3 positions are masked out (spaced seed pattern 1010011). Second is SIMD-accelerated Smith-Waterman of *ν* sequences, implemented using a modified Parasail library (Daily 2016). If the *ν* score is high enough, the reversed query *ν* sequence is aligned to the target *ν* sequence, a weighted sum of the forward and reversed score is calculated; the (query,target) pair is discarded if this sum is below a threshold. Otherwise, the pair is aligned using the *M ^′^*alphabet, and finally the reversed query *M ^′^* profile is aligned to the target if the forward *M ^′^* score is high enough. If all these steps are completed successfully, enough information has accumulated to enable calculation of the test statistic (TS) and *P*-value from log-linear fitting (Edgar and Sahakyan 2025).

### Collection accuracy

“Collection” models a search which is designed to identify as many related structures as possible while allowing an acceptable number of FPs. At a given score cutoff, TPs and FPs are counted in the obvious way, enabling calculation of sensitivity, false-positive errors per query (FPEPQ), precision, recall, true-positive rate (TPR), and false-positive rate (FPR), as needed for CVE, PR and ROC plots. Truth standards family, superfamily and fold are self-explanatory. An additional truth standard, which I call “superfamilyx”, has also been used in the literature, notably in the pioneering paper (Brenner et al. 1998) which introduced homology search validation by SCOP. With superfamilyx, TPs are hits to the the same superfamily, FPs are hits to the same fold, and hits to different superfamilies in the same fold are considered ambiguous and discarded. This ensures that TPs are homologous with high confidence, and naively that FPs are false positives with high confidence. However, SCOP assigns several clearly homologous superfamilies to different folds, and I would therefore argue that these should be discarded as in SCOP40c (Edgar and Sahakyan 2025) so that FPs also have high confidence. A further issue with this standard is that the number of ambiguous hits in an all-vs-all SCOP40 comparison is greater than the number of TPs; discarding them thus ignores a large number of the most challenging cases so that the boundary between TPs and FPs is not sharp. Rather, TPs and FPs are separated by a large gray area, and consequently the task of discriminating TPs from FPs is unrealistically easy. For evaluating sequence search, this issue is often negligible, but structure alignment readily identifies similar structures deep into this gray area. Using homology as a standard is unavoidably problematic for structure, because ultimately it is unknowable whether structures having the same fold but no detectable sequence similarity are explained by homology or by convergent evolution. As an operational solution to this challenge, I regard SCOP family, superfamily and fold categories as gold standards, while noting that there is some unavoidable arbitrariness in all levels of such a classification scheme, and the correspondence with homology cannot be fully known.

Given an accuracy plot, algorithm *A* is unambiguously more accurate than algorithm *B* if all-vs-all scores are available for both algorithms and *A*’s curve is above *B*’s (PR, ROC) or below *B*’s (CVE) over all score cutoffs. However, of the tested algorithms, TM-align is the only one to report all scores. Reseek and Foldseek discard most possible hits via filtering, and DALI does not offer an option to report hits with *Z <* 2. Also, while curves can often be clearly ranked in the higher-scoring regime, they often intersect near the fuzzy boundary between high- and low-scoring hits. Measuring areas over restricted plot ranges can be problematic owing to truncated and step-like curves. Instead, to quantify accuracy and enable algorithm ranking, I designed the following summary metrics. For the CVE curve, I chose to measure sensitivity at three values of `FPEPQ`: 0.1, 1 and 10, corresponding to *E*-value cutoffs 0.1, 1 and 10, respectively, giving `SEPQ0.1` (“sensitivity at `FPEPQ=0.1`”), `SEPQ1` and `SEPQ10`. These cutoffs are higher than *E*-values typically used in practice, but SCOP40 is too small to measure PFEPQ much less than 0.1. The summary metric Sum3 is calculated as the weighted sum `Sum3 = 2*SEPQ0.1 + 1.5*SEPQ1 + SEPQ10`, giving higher weight to lower FPEPQs. From a PR curve, I chose to measure PR90, calculated as recall at precision=90%. Given that CVE, PR and ROC correlate, I chose to focus on `Sum3` and `PR90` rather than introducing any further summary metrics.

### Categorization accuracy

“Categorization” models a search which is designed to classify the query by assigning it to a category which is annotated in the database; for example, the user might wish to classify a structure to a SCOP fold. In practice, a user will typically look at the top few hits. If most or all of the top hits belong to the same fold and have low *E*- or *P*-value, the query can be assigned to the this fold with high confidence. The simplest strategy is to consider the top hit only. A more sophisticated approach which considers other hits would require adding a new algorithm on top of alignment, which would potentially introduce bias and would raise further questions; in particular, would some other design work better? Therefore, I model categorization by considering the top hit only. If there is a hit above the threshold, the top hit is either TP or FP depending on whether it belongs to the same category. If there is no hit, this is considered a TN or FN depending on whether the category exists in the database. Conceptually, collection treats alignment as a binary classification problem (“does this pair belong to the same category, true or false?”) while categorization treats alignment as a multi-class classification problem (“does the query belong to category *m*, *m* = 1*..M* ?”). With a reference of size *N* structures, collection considers *N* ^2^ *− N* tests (all pairs excluding self-hits), while categorization considers *N* tests, one for each query. With collection, leave-one-out challenges a classifier because the bulk of the TP distribution is swamped by FP scores (Edgar and Sahakyan 2025); however, with categorization leave-one-out would create mostly easy tests. Consider classification to superfamily. The top hit will usually belong to the same family, and classification to superfamily is unlikely to fail. To create a challenging test, I leave out other members of the same family. Now the alignment score must distinguish a different family in the same superfamily from the most similar family in a different superfamily. If there is only one family in a given superfamily, no TPs are possible if the rest of the family is held out. Therefore, to achieve correct normalization (e.g., maximum sensitivity = 1), a query is considered only if its superfamily has at least two families. Thus, a (query, target) pair (*q, t*) is considered only if *q* belongs to a superfamily with two or more families, and *t* belongs to a different family from *q*. This enables counting of TPs and FPs based on challenging tests, each of which models a scenario which occurs in practice, i.e. recognition of a category from a distant relative. With this construction, a TP is always possible, so a different strategy is required for measuring TNs and FPs when the category of *q* is absent from the database, and the correct prediction is then that the query does not belong to a known category. This is straightforwardly accomplished by leaving out all other members of the superfamily. Together, I call these methods for calculating a confusion matrix for superfamily categorization the “topsf” standard and the corresponding strategy for fold “topfold”. These standards enable construction of CVE, PR and ROC plots by sweeping over the cutoff, similar to those for collection standards, and a summary statistic `Top3 = 2*SEPQ0.001 + 1.5*SEPQ0.01 + SEPQ0.1`.The error rate thresholds are set to 0.1%, 1% and 10% because these are measurable on a small reference such as SCOP40c, and reflect a plausible range of rates which might be considered acceptable in practice.

## Conclusion

Reseek version 3 substantially improves accuracy in protein structure search, while achieving practical speed and scalability for multi-million-structure databases. Training of separate statistical models reveals that distinct features are informative for family, superfamily and fold relationships. These results establish that Reseek advances the state of the art for search of AI-scale protein structure databases.

## Code availability

https://github.com/rcedgar/reseek

