## Supplementary information for "Rich structure alphabets enable highest accuracy protein search"

Robert C. Edgar

### Supplementary Information

### Supplementary Figures

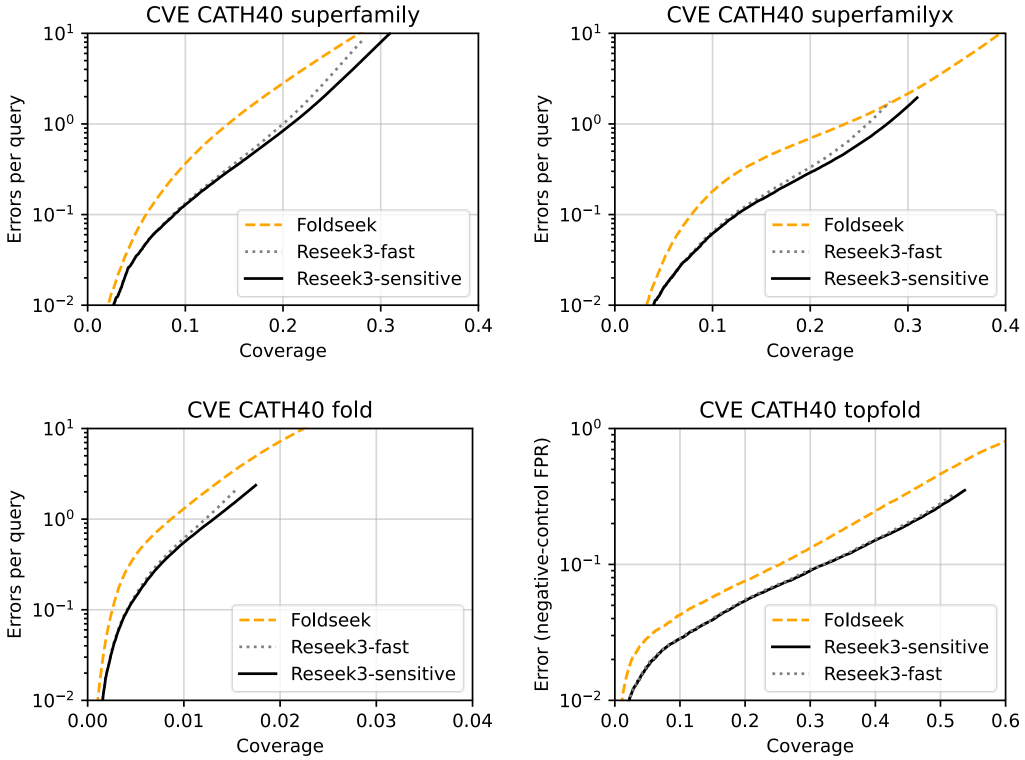


Supp Fig. S1. CVE comparison of Foldseek and Reseek on CATH40 reference.

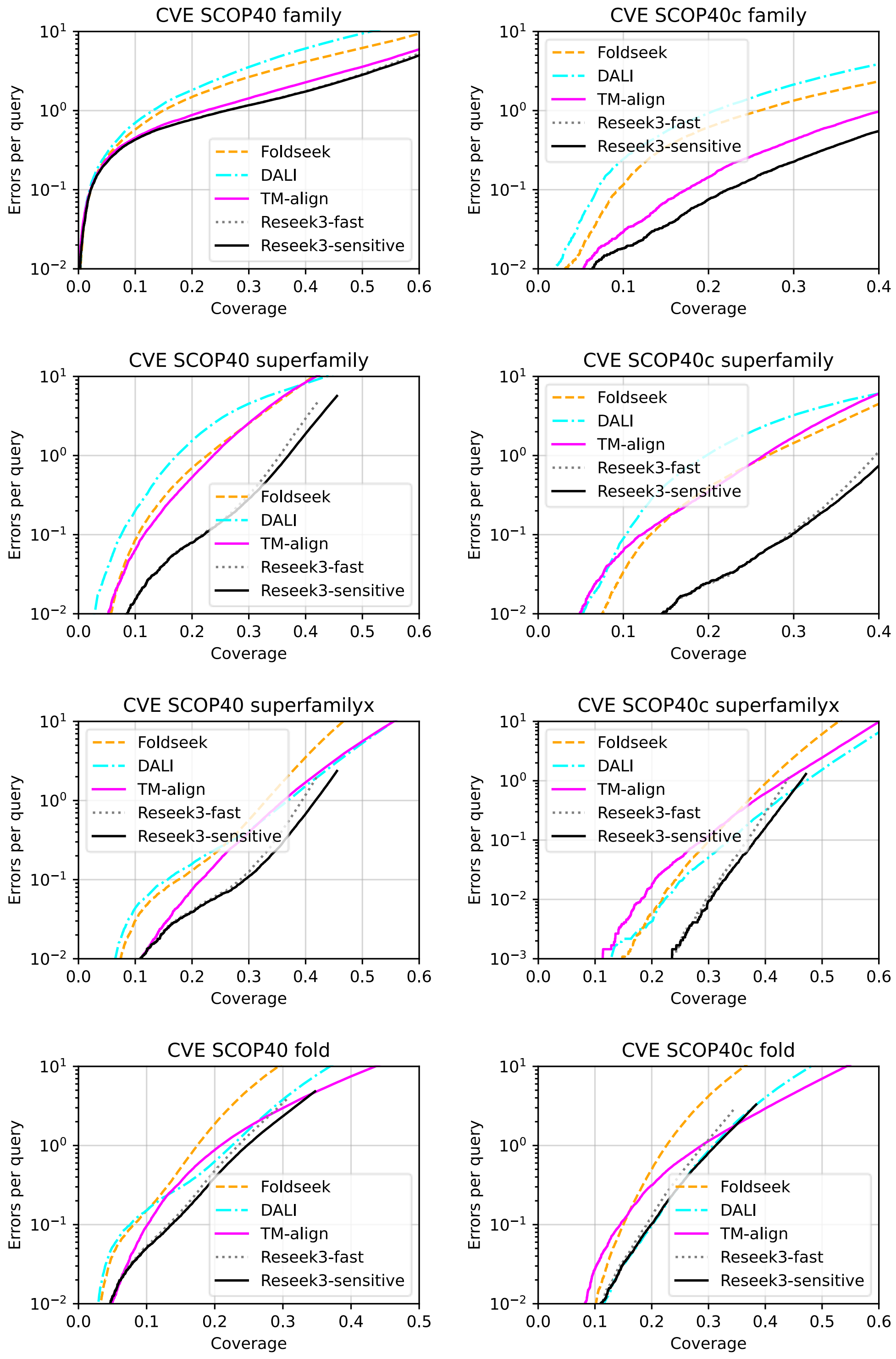


Supp Fig. S2. CVE comparison of methods on SCOP40 and SCOP40c references.

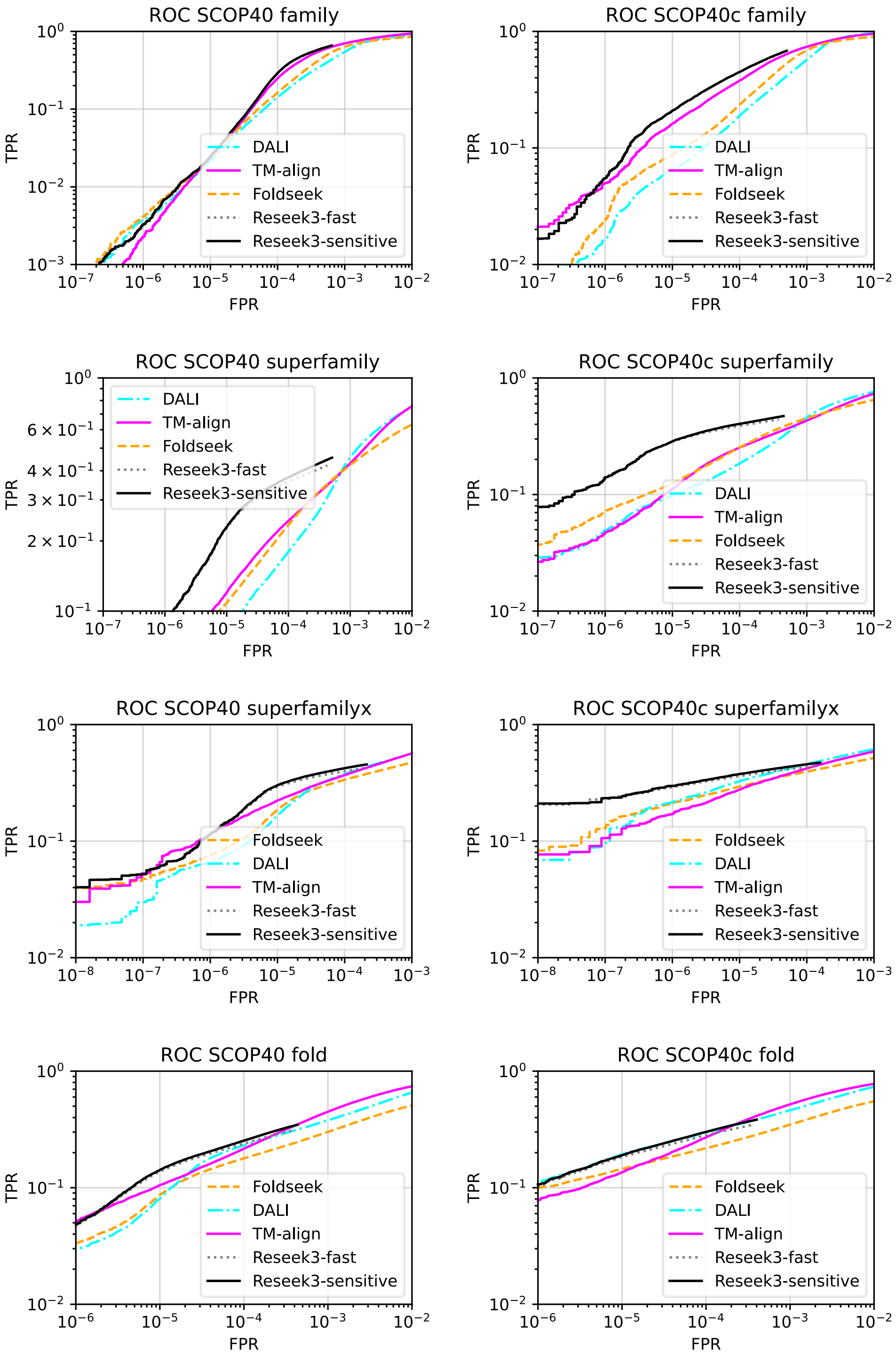


Supp Fig. S3 ROC comparison of methods on SCOP40 and SCOP40c references using family, superfamily, superfamilyx and fold standards.

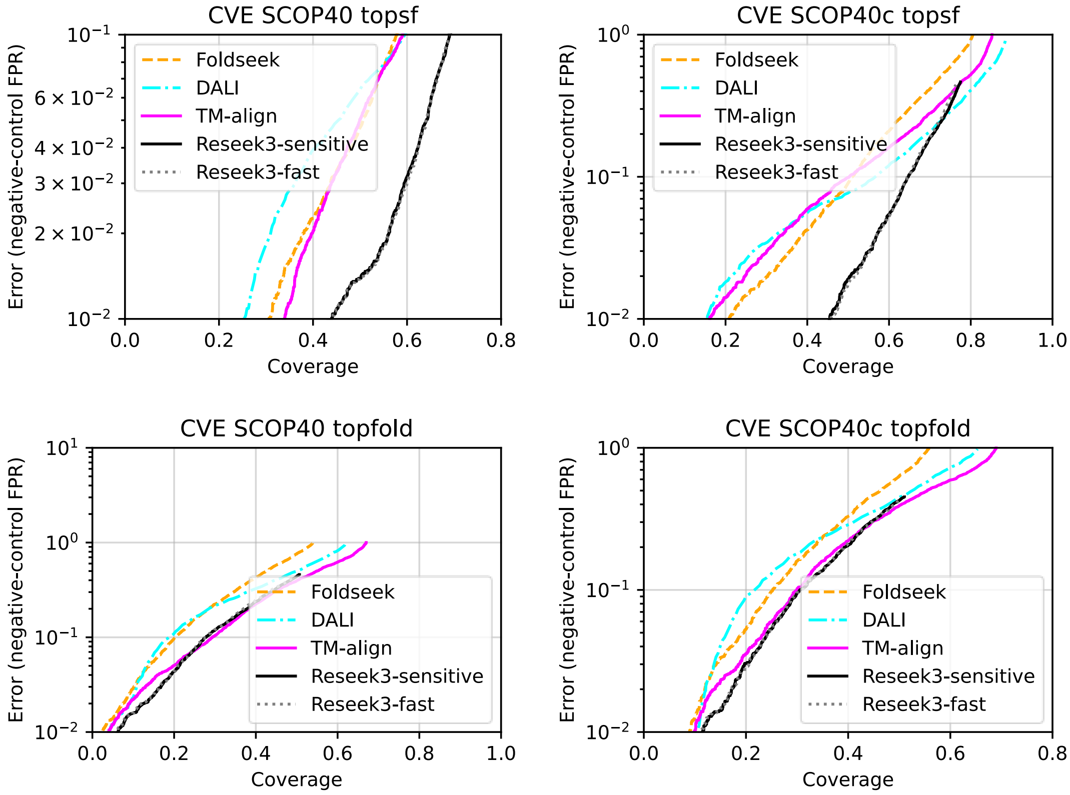


Supp Fig. S4 CVE method comparison using SCOP40 and SCOP40c references using topsf and topfold standards.

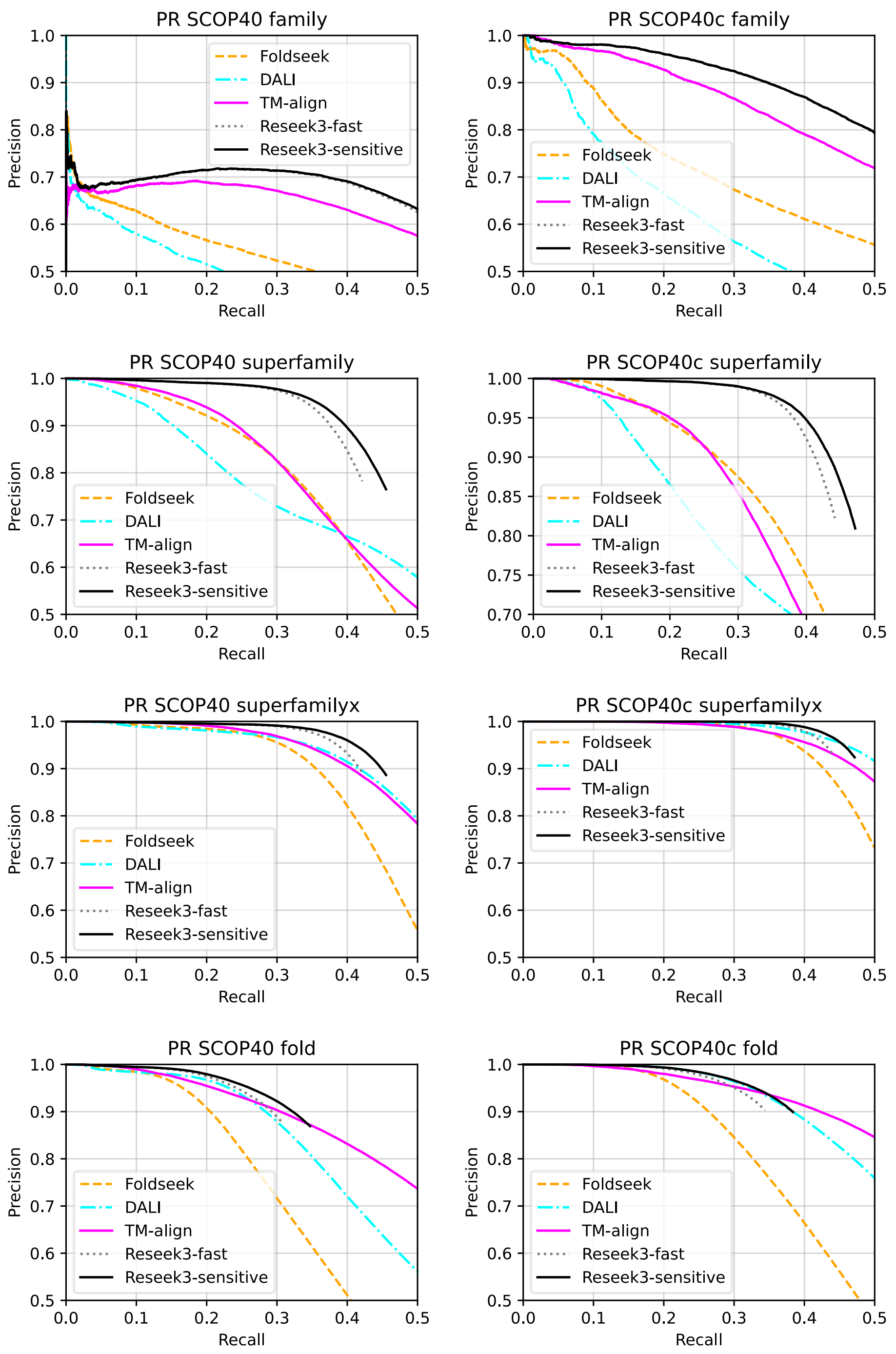


Supp Fig. S4 PR comparison of methods on SCOP40 and SCOP40c references using family, superfamily, superfamilyx and fold standards.

### Supplementary Data -- Metric values

Sum3>family>cath40

| SEPQ0.1 | SEPQ1 | SEPQ10 | Sum3 | algo |
|---|---|---|---|---|
| 0.028 | 0.117 | 0.188 | 0.420 | reseek3_fast |
| 0.028 | 0.117 | 0.202 | 0.434 | reseek3_sensitive |
| 0.060 | 0.144 | 0.278 | 0.613 | foldseek |

Sum3>family>scop40

| SEPQ0.1 | SEPQ1 | SEPQ10 | Sum3 | algo |
|---|---|---|---|---|
| 0.019 | 0.130 | 0.507 | 0.741 | dali |
| 0.022 | 0.149 | 0.613 | 0.882 | foldseek |
| 0.022 | 0.259 | 0.630 | 1.062 | reseek3_fast |
| 0.022 | 0.224 | 0.685 | 1.064 | tm |
| 0.022 | 0.259 | 0.654 | 1.087 | reseek3_sensitive |

Sum3>family>scop40x

| SEPQ0.1 | SEPQ1 | SEPQ10 | Sum3 | algo |
|---|---|---|---|---|
| 0.068 | 0.207 | 0.627 | 1.074 | dali |
| 0.094 | 0.260 | 0.725 | 1.303 | foldseek |
| 0.174 | 0.404 | 0.762 | 1.716 | tm |
| 0.222 | 0.475 | 0.663 | 1.820 | reseek3_fast |
| 0.222 | 0.475 | 0.681 | 1.838 | reseek3_sensitive |

Sum3>fold>cath40

| SEPQ0.1 | SEPQ1 | SEPQ10 | Sum3 | algo |
|---|---|---|---|---|
| 0.003 | 0.009 | 0.022 | 0.041 | foldseek |
| 0.004 | 0.012 | 0.015 | 0.042 | reseek3_fast |
| 0.004 | 0.013 | 0.017 | 0.045 | reseek3_sensitive |

Sum3>fold>scop40

| SEPQ0.1 | SEPQ1 | SEPQ10 | Sum3 | algo |
|---|---|---|---|---|
| 0.081 | 0.174 | 0.293 | 0.718 | foldseek |
| 0.075 | 0.223 | 0.364 | 0.850 | dali |
| 0.132 | 0.235 | 0.309 | 0.925 | reseek3_fast |
| 0.101 | 0.208 | 0.436 | 0.951 | tm |
| 0.136 | 0.247 | 0.347 | 0.991 | reseek3_sensitive |

Sum3>fold>scop40x

| SEPQ0.1 | SEPQ1 | SEPQ10 | Sum3 | algo |
|---|---|---|---|---|
| 0.150 | 0.226 | 0.363 | 1.003 | foldseek |

```
  0.191  0.291   0.344  1.163  reseek3_fast
  0.199  0.313   0.384  1.251  reseek3_sensitive
  0.145  0.288   0.543  1.264  tm
  0.199  0.303   0.478  1.330  dali

Sum3>superfamily>cath40
SEPQ0.1  SEPQ1  SEPQ10  Sum3   algo
  0.060  0.144   0.278  0.613  foldseek
  0.087  0.200   0.282  0.756  reseek3_fast
  0.089  0.209   0.309  0.800  reseek3_sensitive

Sum3>superfamily>scop40
SEPQ0.1  SEPQ1  SEPQ10  Sum3   algo
  0.075  0.171   0.432  0.837  dali
  0.105  0.227   0.413  0.963  foldseek
  0.115  0.237   0.418  1.004  tm
  0.218  0.354   0.421  1.388  reseek3_fast
  0.221  0.369   0.454  1.450  reseek3_sensitive

Sum3>superfamily>scop40x
SEPQ0.1  SEPQ1  SEPQ10  Sum3   algo
  0.102  0.193   0.493  0.987  dali
  0.122  0.265   0.452  1.092  tm
  0.132  0.269   0.469  1.136  foldseek
  0.294  0.396   0.442  1.624  reseek3_fast
  0.298  0.413   0.471  1.687  reseek3_sensitive

Sum3>superfamilyx>cath40
SEPQ0.1  SEPQ1  SEPQ10  Sum3   algo
  0.078  0.236   0.395  0.905  foldseek
  0.121  0.259   0.282  0.912  reseek3_fast
  0.125  0.279   0.309  0.977  reseek3_sensitive

Sum3>superfamilyx>scop40
SEPQ0.1  SEPQ1  SEPQ10  Sum3   algo
  0.176  0.330   0.466  1.314  foldseek
  0.156  0.366   0.549  1.409  dali
  0.216  0.361   0.555  1.529  tm
  0.285  0.393   0.421  1.580  reseek3_fast
  0.293  0.417   0.454  1.665  reseek3_sensitive

Sum3>superfamilyx>scop40x
```

| SEPQ0.1 | SEPQ1 | SEPQ10 | Sum3 | algo |
|---|---|---|---|---|
| 0.303 | 0.405 | 0.530 | 1.743 | foldseek |
| 0.290 | 0.434 | 0.601 | 1.832 | tm |
| 0.368 | 0.438 | 0.442 | 1.835 | reseek3_fast |
| 0.382 | 0.463 | 0.471 | 1.931 | reseek3_sensitive |
| 0.334 | 0.466 | 0.630 | 1.997 | dali |

PR90>family>cath40

| PR90 | algo |
|---|---|
| 0.1729 | reseek3_fast |
| 0.179 | reseek3_sensitive |
| 0.2013 | foldseek |

PR90>family>scop40x

| PR90 | algo |
|---|---|
| 0.05771 | dali |
| 0.09115 | foldseek |
| 0.2449 | tm |
| 0.3472 | reseek3_fast |
| 0.3475 | reseek3_sensitive |

PR90>fold>cath40

| PR90 | algo |
|---|---|
| 0.01542 | reseek3_fast |
| 0.01746 | reseek3_sensitive |
| 0.02242 | foldseek |

PR90>fold>scop40

| PR90 | algo |
|---|---|
| 0.2051 | foldseek |
| 0.281 | dali |
| 0.2924 | reseek3_fast |
| 0.3047 | tm |
| 0.3221 | reseek3_sensitive |

PR90>fold>scop40x

| PR90 | algo |
|---|---|
| 0.2639 | foldseek |
| 0.3442 | reseek3_fast |
| 0.3806 | dali |
| 0.3832 | reseek3_sensitive |
| 0.4221 | tm |

PR90>superfamily>cath40

| PR90 | algo |
|---|---|
| 0.2013 | foldseek |
| 0.2507 | reseek3_fast |
| 0.2702 | reseek3_sensitive |

PR90>superfamily>scop40

| PR90 | algo |
|---|---|
| 0.1503 | dali |
| 0.2288 | foldseek |
| 0.2429 | tm |
| 0.3758 | reseek3_fast |
| 0.3969 | reseek3_sensitive |

PR90>superfamily>scop40x

| PR90 | algo |
|---|---|
| 0.1668 | dali |
| 0.2636 | tm |
| 0.2694 | foldseek |
| 0.4128 | reseek3_fast |
| 0.4341 | reseek3_sensitive |

PR90>superfamilyx>cath40

| PR90 | algo |
|---|---|
| 0.2817 | reseek3_fast |
| 0.3094 | reseek3_sensitive |
| 0.3558 | foldseek |

PR90>superfamilyx>scop40

| PR90 | algo |
|---|---|
| 0.3556 | foldseek |
| 0.406 | tm |
| 0.4091 | dali |
| 0.4172 | reseek3_fast |
| 0.4476 | reseek3_sensitive |

PR90>superfamilyx>scop40x

| PR90 | algo |
|---|---|
| 0.4275 | foldseek |
| 0.4416 | reseek3_fast |
| 0.4715 | reseek3_sensitive |

| | |
|---|---|
| 0.4757 | tm |
| 0.5126 | dali |

Top3>topfold>cath40

| TEPQ0.001 | TEPQ0.01 | TEPQ0.1 | Top3 | algo |
|---|---|---|---|---|
| 0.002 | 0.011 | 0.254 | 0.274 | foldseek |
| 0.003 | 0.022 | 0.320 | 0.358 | reseek3_fast |
| 0.003 | 0.022 | 0.322 | 0.361 | reseek3_sensitive |

Top3>topfold>scop40

| TEPQ0.001 | TEPQ0.01 | TEPQ0.1 | Top3 | algo |
|---|---|---|---|---|
| 0.003 | 0.020 | 0.202 | 0.237 | foldseek |
| 0.005 | 0.033 | 0.191 | 0.250 | dali |
| 0.005 | 0.039 | 0.294 | 0.363 | tm |
| 0.005 | 0.060 | 0.277 | 0.376 | reseek3_fast |
| 0.005 | 0.062 | 0.278 | 0.380 | reseek3_sensitive |

Top3>topfold>scop40x

| TEPQ0.001 | TEPQ0.01 | TEPQ0.1 | Top3 | algo |
|---|---|---|---|---|
| 0.030 | 0.106 | 0.219 | 0.437 | dali |
| 0.032 | 0.089 | 0.253 | 0.451 | foldseek |
| 0.014 | 0.098 | 0.299 | 0.475 | tm |
| 0.045 | 0.112 | 0.310 | 0.568 | reseek3_fast |
| 0.047 | 0.114 | 0.306 | 0.571 | reseek3_sensitive |

Top3>topsf>scop40

| TEPQ0.001 | TEPQ0.01 | TEPQ0.1 | Top3 | algo |
|---|---|---|---|---|
| 0.083 | 0.255 | 0.597 | 1.146 | dali |
| 0.162 | 0.307 | 0.578 | 1.362 | foldseek |
| 0.176 | 0.340 | 0.591 | 1.452 | tm |
| 0.192 | 0.440 | 0.689 | 1.732 | reseek3_fast |
| 0.192 | 0.439 | 0.690 | 1.734 | reseek3_sensitive |

Top3>topsf>scop40x

| TEPQ0.001 | TEPQ0.01 | TEPQ0.1 | Top3 | algo |
|---|---|---|---|---|
| 0.019 | 0.161 | 0.503 | 0.783 | tm |
| 0.067 | 0.156 | 0.568 | 0.937 | dali |
| 0.096 | 0.208 | 0.510 | 1.014 | foldseek |
| 0.194 | 0.455 | 0.647 | 1.717 | reseek3_sensitive |
| 0.202 | 0.458 | 0.645 | 1.735 | reseek3_fast |
